# DynaRNA: Dynamic RNA Conformation Ensemble Generation with Diffusion Model

**DOI:** 10.1101/2025.05.22.655453

**Authors:** Zhengxin Li, Junjie Zhu, Xiaokun Hong, Zhuoqi Zheng, Taeyoung Cui, Yutong Sun, Ting Wei, Haifeng Chen

**Author notes:** Corresponding Author, Hai-Feng Chen (Full Professor), State Key Laboratory of Microbial metabolism, Joint International Research Laboratory of Metabolic & Developmental Sciences, Department of Bioinformatics and Biostatistics, National Experimental Teaching Center for Life Sciences and Biotechnology, School of Life Sciences and Biotechnology, Shanghai Jiao Tong University, Shanghai, 200240, China; **Email:**. Notes The authors declare that there is no conflict of interest.

## Abstract

RNA plays a wide variety of roles in biological processes. In addition to serving as the coding messenger RNA (mRNA), the vast majority of RNAs function as non-coding RNAs (ncRNAs), where their dynamic structural ensemble is critical for mediating diverse biological functions. However, traditional experimental techniques and molecular dynamics (MD) simulations face significant challenges in characterizing the conformational dynamics of RNA, due to inherent methodological limitations and high computing power cost. We herein presented DynaRNA, a diffusion-based generative model for RNA conformation ensemble. DynaRNA employs denoising diffusion probabilistic model (DDPM) with equivariant graph neural network (EGNN) to directly model RNA 3D coordinates, enabling rapid exploration of RNA conformational space. DynaRNA enables end-to-end generation of RNA conformation ensemble reproducing experimental geometries without the need for Multiple Sequence Alignments (MSA) information. Our results demonstrate that DynaRNA effectively and accurately generate tetranucleotides ensemble with lower intercalation rate than molecular dynamics simulations. Besides, DynaRNA has the ability to capture rare excited states of HIV TAR(trans-activation response) element, and recapitulate de novo folding of tetraloops. DynaRNA serve as a versatile and efficient platform for modeling RNA structural dynamics, with broad implications potential in RNA structural biology, synthetic biology, and therapeutic development.

## Introduction

Over 95% of human genome is transcribed to non-coding RNA, which serve pivotal roles in biomolecular processes.^1^ The intrinsic dynamic flexibility and pronounced conformational heterogeneity of RNA endow it with diverse functional capabilities.^2^ Deciphering the conformational ensembles of RNA is fundamental for understanding its intricate mechanisms of action, advancing RNA-targeted drug discovery, and facilitating the design of RNA-based therapeutic strategies.^3^ However, traditional experimental methods including NMR, X-ray, and cryo-electron microscopy encounter considerable limitations in resolving the complex conformational ensembles of RNA.^4^ On the one hand, these methods often average signals from multiple conformations, making it difficult to accurately capture RNA’s highly heterogeneous structural characteristics.^5^ On the other hand, the intrinsic properties of RNA structures further complicate their resolution by experimental approaches.^6^ Conventional computational methods such as Molecular dynamics simulations (MDs) is too expensive to explore RNA’s vast conformation space.^7^ Besides, inaccuracies in RNA force fields severely limit the application of MDs.^8–11^

Recently, the rapid advancement of artificial intelligence methodologies has provided novel opportunities for structural biology.^12^ AlphaFold2 significantly improved the accuracy of protein structure prediction but does not include RNA.^13^ AlphaFold3 introduced a substantially updated diffusion-based architecture which extend beyond protein to nucleic acids and other biomolecule structure prediction.^14^ AlphaFold3 is predominantly confined to predicting single stable conformations rather than generating conformation ensemble of RNA, which is important for comprehensively characterizing the heterogeneity of RNA.^15^ Diffusion model has been shown promise in generating protein conformation ensemble^16^, but has not been used in predicting RNA conformation ensemble to our best knowledge.

In this work, we developed DynaRNA to generate RNA dynamic conformation ensemble based on a particular generative model. This approach represents the first attempt of application of diffusion model directly modeling 3D coordinates of RNA for conformation ensemble generation. We herein demonstrated the capability of diffusion model in RNA conformation generation with developing DynaRNA model in an attempt for directly modeling 3D coordinates of RNA, orders of magnitude faster than MDs. We show that DynaRNA can generalize across various molecular systems and propose diverse structures that agreement with experimental results. We employed several RNA molecular systems, including tetranucleotides^17^, tetraloop^18–19^, HIV TAR^20–21^ to demonstrate applications of DynaRNA. DynaRNA showed the ability to generate RNA dynamic conformation ensemble in agreement with experimental observations. Besides, DynaRNA can capture the excited-ground state of HIV TAR and de novo folding of tetraloop. These results demonstrate that DynaRNA extends the application of diffusion models to RNA conformational ensemble generation, offering a novel approach for exploring vast dynamic conformational space of RNA.

## Results

### DynaRNA architecture

Diffusion models have been widely used and proven effective in molecule generation. In this study, we employed a denoising diffusion probabilistic model (DDPM) tailored for RNA conformational ensemble generation, which operates directly on the 3D atomic coordinates of a given input structure. Distinct from conventional DDPMs that gradually diffuse inputs into pure Gaussian noise^22^, our model adopts a partial noising scheme, where the diffusion is applied only up to an intermediate noise step rather than a full corruption. This enables a tunable balance between preserving structural information from the original input and introducing stochastic variability for sampling diverse conformations.^23^ The generative pipeline consists of two stages: a forward diffusion process that incrementally adds Gaussian noise to the coordinate space, and a reverse denoising process that iteratively reconstructs the conformation. The denoising network is implemented using Equivariant Graph Neural Networks (EGNNs)^24^, which are designed to respect the Euclidean symmetries (E(3)) of molecular structures, such as rotation and translation equivariance. By modeling the molecule as a spatial graph, EGNNs ensure that the generative process remains consistent with the physical geometry of the system.^25^ The flexibility introduced by partial noising allows users to control how much structural prior is retained during sampling, enabling tailored conformational generation based on the desired balance between fidelity and diversity. This makes our approach particularly suitable for modeling RNA molecules, as RNA conformation ensembles are often more diverse than protein. Further architectural and training details are provided in the Materials and Methods section.

### General Validation of DynaRNA

We first assessed the geometric fidelity of RNA conformations generated by DynaRNA by examining two key structural features: the distance between adjacent nucleotides and the hyper bond angles formed by three consecutive nucleotide C4’ atoms. These metrics serve as important indicators of RNA backbone integrity. We computed the distributions of these features in the DynaRNA-generated conformations and compared them with reference distributions derived from high-resolution RNA structures in the Protein Data Bank (PDB)^26^, as shown in Figure 2. The results demonstrated that the adjacent C4’–C4’ distances in the DynaRNA-generated ensemble are highly consistent with those observed in experimental structures, both peaking around 6 Å. Besides, the hyper bond angles defined by three consecutive C4’ atoms are centered around 40 degrees in both datasets, reinforcing the model’s ability to reproduce the local backbone geometry of native RNA conformations. Taken together, these observations suggest that DynaRNA is capable of generating RNA conformations with high geometric plausibility, closely matching the statistical features of experimentally resolved RNA structures. This level of agreement underscores the model’s fidelity and its potential utility in RNA structure modeling and related computational studies.

**Figure 1.**
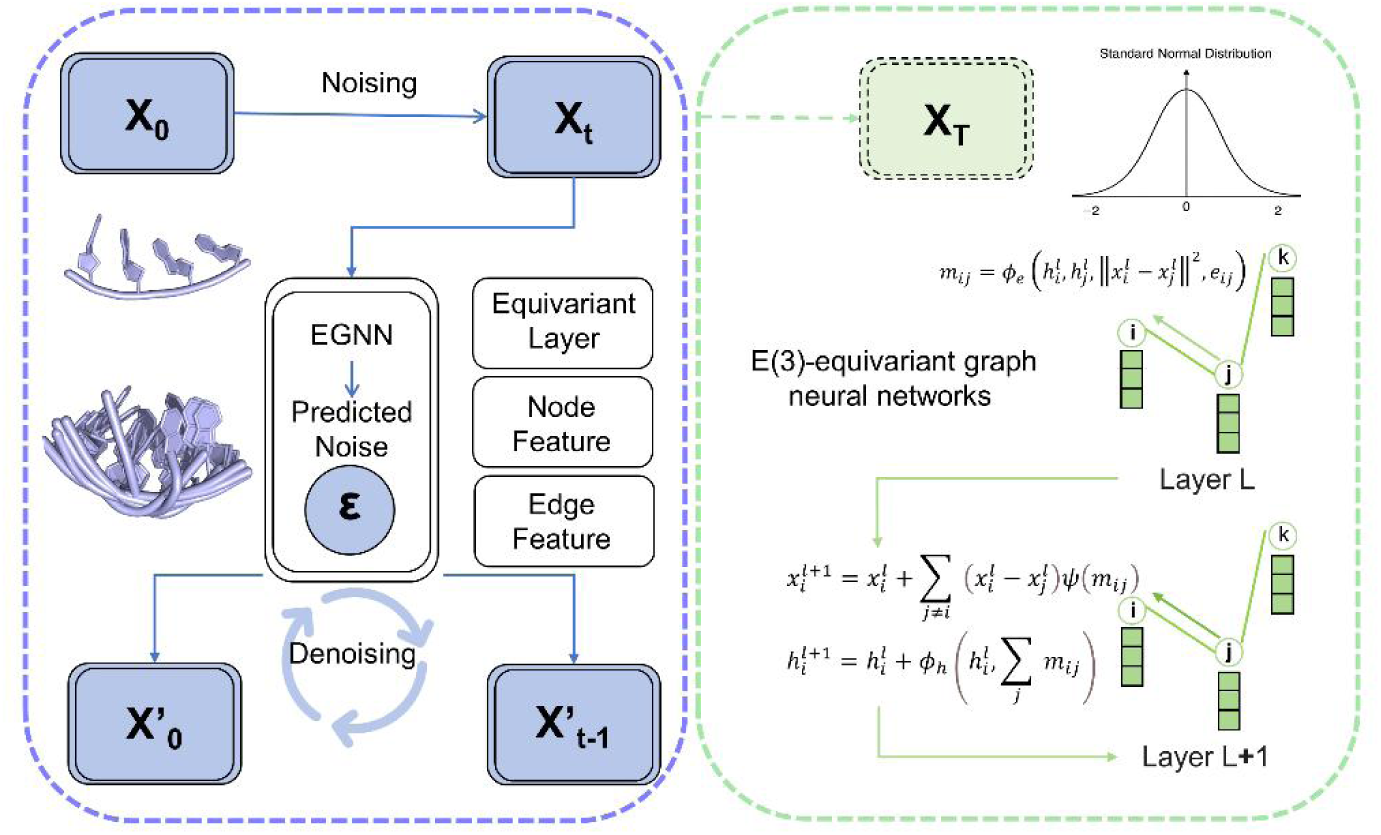
Overview of DynaRNA architecture. The framework comprises two processes: a forward noising process, in which Gaussian noise is progressively added to the input structure, and a reverse denoising process, wherein Equivariant Graph Neural Networks (EGNNs) are utilized to predict the noise and denoising.

**Figure 2.**
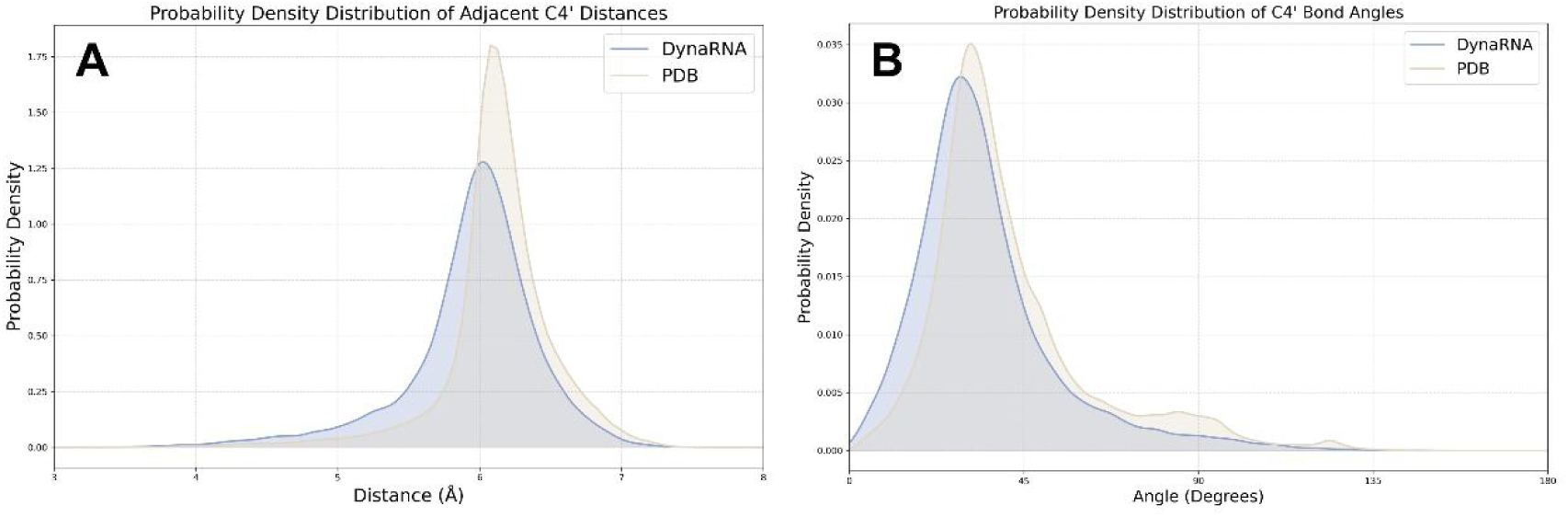
Fidelity validation of DynaRNA-generated conformation ensemble. A. Distribution of adjacent of C4’ distances of conformation ensembles generated by DynaRNA(blue) compared to PDB experimental structures(orange). B. Distribution of C4’ hyper bond angles of conformation ensembles generated by DynaRNA(blue) compared to PDB experimental structures(orange).

### DynaRNA can capture the conformation ensemble of tetranucleotides

Tetranucleotide, consisting of four nucleotides, serve as a key benchmark system for RNA computational structure research.^5^ Existing computational methods, such as molecular dynamics simulations, generate a large number of RNA intercalated conformations, which are in serious disagreement with the results of solution NMR experiments.^17^ We systematically compared the performance of DynaRNA with MD simulations employing three distinct force fields (OL3^27^, BSFF1^11^, BSFF2^10^) initialized from canonical A-form structures and intercalated conformations. Quantitative analysis of intercalation propensity in the generated conformational ensembles is presented in Figure 3. Notably, DynaRNA-predicted tetranucleotide ensembles exhibited substantially lower intercalation ratios compared to MD simulations with OL3. This improvement was particularly evident when starting from the intercalated conformation. For CAAU and CCCC systems, simulations with OL3 became trapped in the intercalated conformation, with intercalation rates of 97.3% and 90.7% respectively. In contrast, DynaRNA yielded conformation ensemble with intercalation rates of only 9.2% and 4.7%. Regardless of whether the simulations started from the A-form or the intercalated conformation, the intercalation rates in DynaRNA-generated ensembles remained below 10%, effectively demonstrating the robustness of DynaRNA.

**Figure 3.**
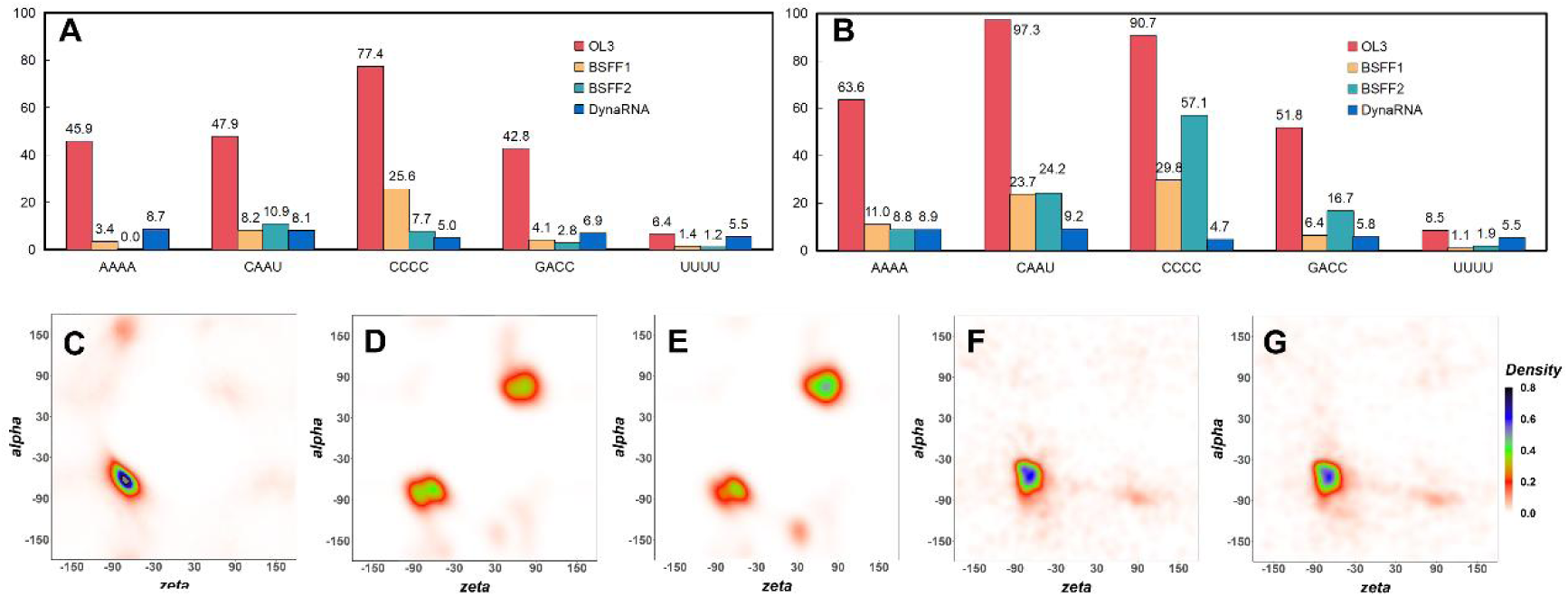
DynaRNA captures the experimental conformation ensemble of tetranucleotides. A. Intercalation ration of conformation ensembles generated with molecular dynamics simulations and DynaRNA initialized from A-form structures. B. Intercalation ration of conformation ensembles generated with molecular dynamics simulations and DynaRNA initialized from intercalated structures. C. Dihedral statistical distribution of PDB experimental structures. D. Dihedral statistical distribution of conformation ensemble generated with molecular dynamics simulations with OL3 initialized from A-form structures. E. Dihedral statistical distribution of conformation ensemble generated with molecular dynamics simulations with OL3 initialized from intercalated structures. F. Dihedral statistical distribution of conformation ensemble generated with DynaRNA initialized from A-form structures. G. Dihedral statistical distribution of conformation ensemble generated with DynaRNA initialized from intercalated structures.

Furthermore, we performed detailed structural analysis of the conformational ensembles. Previous studies have shown significant differences in the ζ/α dihedral distributions between intercalated and non-intercalated conformations.^11^ Our results shown in Fig3 revealed that in the ensemble simulated with OL3, the ζ/α dihedrals were predominantly concentrated in the intercalated conformation region (+30° to +90°). In contrast, DynaRNA-generated ensembles exhibited ζ/α dihedrals concentrated in the non-intercalated region (−30° to −90°). Structural clustering analysis shown in Figure S1 and S2 further confirmed that the major conformation in DynaRNA-generated ensembles was the non-intercalated A-form, whereas OL3 force field simulations predominantly yielded intercalated conformations. Besides, we also analyzed the dihedral distribution of conformation ensemble generated by molecular dynamics simulations with BSFF1 and BSFF2. Results shown in Fig S3-S6 demonstrated that DynaRNA attains accuracy on par with, or exceeding, that of MD simulations in RNA conformation ensemble generation, while also offering significantly faster computational speed due to its innovative approach of bypassing the need for step-by-step sampling.

### DynaRNA can capture the excited states of RNA conformation

The HIV-1 Trans-Activation Response Element (TAR) has emerged as a highly promising therapeutic target.^28^ Its structure consisting of two helical regions connected by a bulge and a hairpin loop motif at the apex has attracted extensive research attention. ^29^Previous studies have revealed that besides the dominating ground state (GS), HIV-1 TAR also adopts low-populated excited states (ES).^20–21^ These ES play essential roles in biochemical reactions, disease mechanisms, and therapeutic development. ^30^However, due to their richness in non-canonical mismatches and energetically unfavorable nature, they are sparse and short-lived, posing significant challenges for structural characterization. Conventional experimental techniques struggle to capture RNA excited states.^31^ Traditional computational techniques such as molecular dynamics simulations face difficulties in overcoming the high energy barriers required to sample ES.^32^ DynaRNA provide the unique technique to overcome the energy barrier and directly explore the ensemble of conformational space. Recently, Ainan Geng et al determined an HIV TAR ES termed ES2 with the population of about 0.4% and a lifetime of ∼2.1 ms.^21^ We herein generated the HIV TAR conformation ensemble with DynaRNA initialized from both GS and ES2. As shown in Fig4, GS differed from ES2 of secondary structure, involving six base pairings and over fifteen nucleotides. Transitions between these states necessitate crossing multiple conformational potential energy barriers, presenting a formidable computational challenge for molecular dynamics simulations. This challenge is particularly serious for ES2, which exists at a higher potential energy level, making it especially difficult to access from GS. Nevertheless, DynaRNA has successfully bridged this gap. When initiated from GS, DynaRNA generated conformational ensembles that effectively captured ES2. Conversely, when initiated from ES2, DynaRNA also successfully sampled GS. This bidirectional capability of DynaRNA is showcased in Fig. 4.

**Figure 4.**
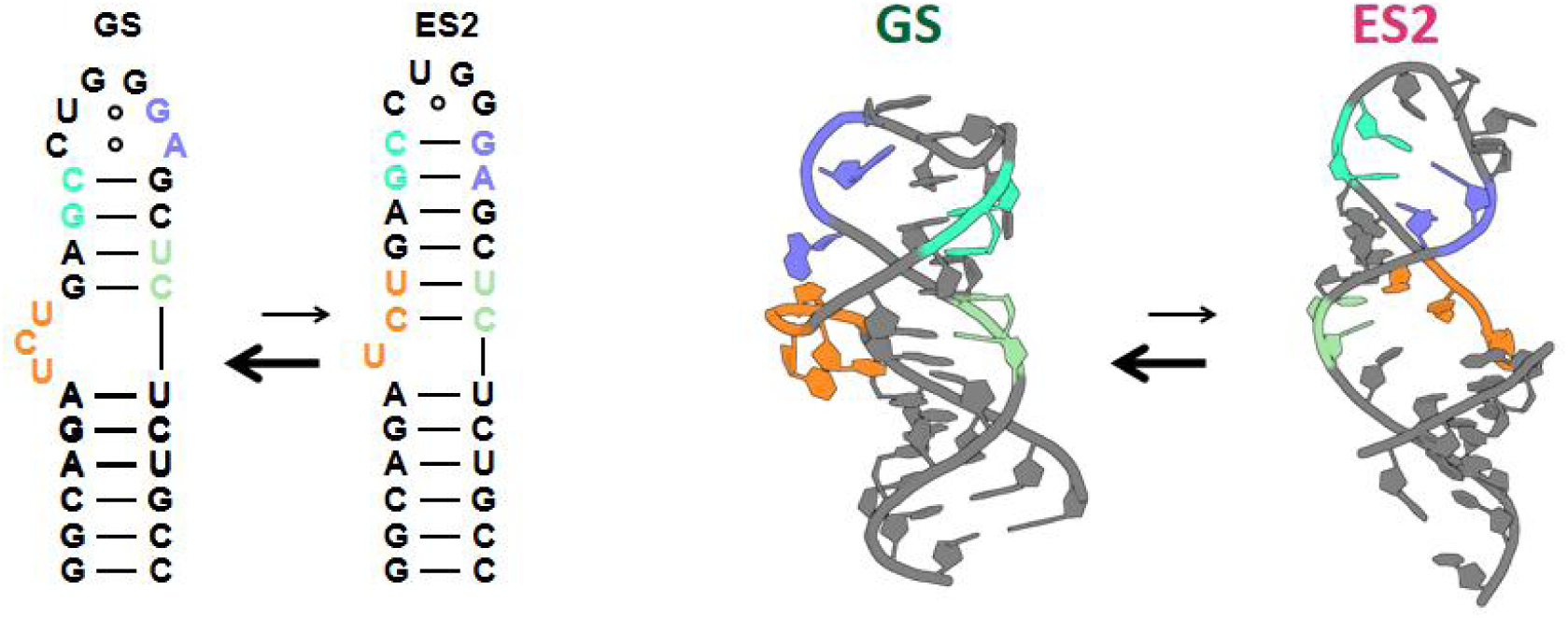
Secondary and tertiary structure of HIV TAR GS and ES2 generated by DynaRNA initialized from the other state.

### DynaRNA can capture de novo folding of RNA tetraloops

Tetraloops, comprising a Watson–Crick base-paired stem and a loop of four nucleotides, represent one of the most ubiquitous and well-characterized RNA secondary structure motifs.^33^ Tetraloops play critical roles in RNA folding, stability, and function, and often serve as nucleation sites in tertiary interactions.^34^ Previous studies^35^ have shown that RNA hairpin folding is a hierarchical process, which makes it computationally expensive to de novo capture the tetraloop folded state using molecular dynamics simulations.^36^ This challenge is further exacerbated by the inaccuracies of current RNA force fields.^37^ Despite these challenges, DynaRNA successfully generated native-like folded conformations of tetraloops starting from fully extended, single-stranded RNA sequences without any structural restraints or prior knowledge. Results of alignment between experimental structure and DynaRNA-predicted structure is shown in Figure 5. DynaRNA achieved the atom root-mean-square deviation (RMSD) of 0.9 Å for the UUCG tetraloop (PDB ID: 2KOC) and 1.3 Å for the GAAA tetraloop (PDB ID: 8CLR) compared with the corresponding native structures, which recapitulated all of the Watson-Crick base pairs with no prior knowledge. Compared to traditional molecular dynamics simulations that require extensive sampling, DynaRNA offers a more efficient and flexible approach for modeling RNA with ability to accurately reconstruct native-like tetraloop structures from scratch, without relying on predefined knowledge or exclusive sampling.

**Figure 5.**
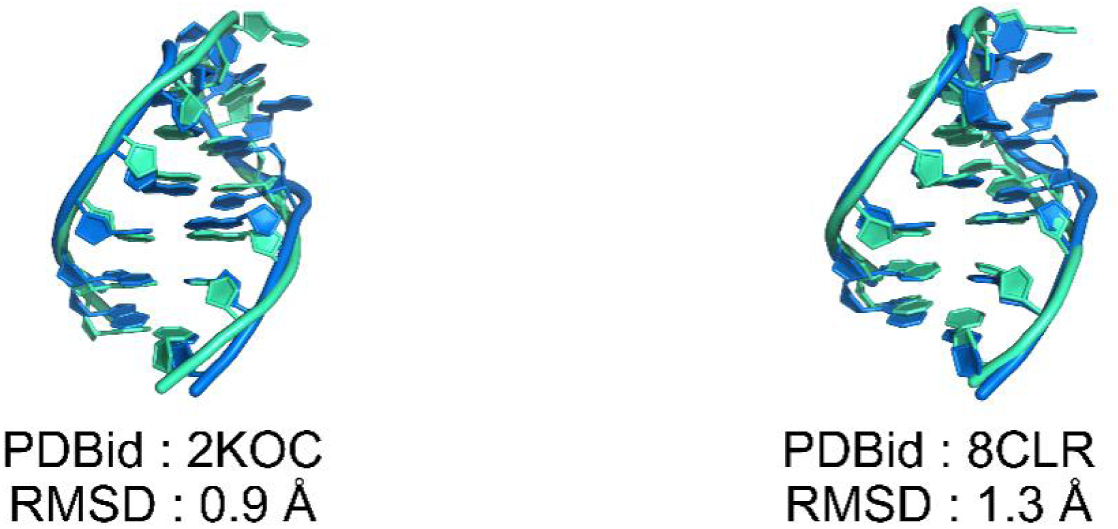
Results of alignment of experimental structure (green) and conformation generated with DynaRNA (blue).

## Discussion

Deciphering the complex hierarchical structural dynamics of RNA is crucial for understanding its functional mechanisms^38^, but this remains highly challenging for both traditional experimental and computational approaches.^39^ To bridge this gap, we employed neural generator to directly sample RNA dynamic conformation (DynaRNA). DynaRNA represents a paradigm shift in computational modeling of RNA conformational dynamics by leveraging the power of diffusion-based generative models. Unlike traditional molecular dynamics simulations that rely on step-by-step sampling with physics-based force fields and are often limited by sampling inefficiencies and force field inaccuracies, DynaRNA efficiently generate diverse and physically plausible RNA conformations.

DynaRNA successfully reproduced experimental geometries (e.g., C4’–C4’ distances, hyper bond angles) and predicted folded tetraloop structure de novo from fully unfolded initial conformation. Besides, DynaRNA outperformed MD simulations in generating tetranucleotide conformation ensembles, achieving intercalation rates below 10% compared to >90% for OL3 force fields and demonstrated orders-of-magnitude faster computational efficiency. While conventional MD simulations typically require weeks of intensive sampling to explore conformational landscapes (often hindered by energy barriers and force field inaccuracies), DynaRNA generates physically plausible ensembles in mere minutes to hours on a single GPU. Notably, DynaRNA achieved high efficiency in capturing rare excited states, capturing the HIV TAR’s low-population (∼0.3%) conformation, highlighting its robustness in escaping energy traps—a critical limitation of physics-based simulations. These results underscored the ability of DynaRNA to accurately and efficiently resolve RNA’s intrinsic heterogeneity.

Furthermore, the framework of DynaRNA holds substantial potential for expansion through three strategic avenues. First, the artificial intelligence generative network of RNA can be integrated with physics-based model (e.g. force field). Refining generated conformations with short MD simulations for local energy minimization could reconcile data-driven efficiency with physical realism. Second, incorporating richer training data, including RNA molecular dynamics trajectories and multi-resolution atomic representations (e.g., explicit backbone atoms beyond C4’), would enhance the model’s ability to capture subtle conformational nuances. Third, the framework of DynaRNA can be expanded to model DNA dynamics^40^, enabling comparative studies of nucleic acid flexibility.

DynaRNA bridges critical gaps in RNA research by complementing both experimental and computational techniques, accelerating RNA therapeutic development, and expanding the scope of AI-driven structural analysis. DynaRNA can combine traditional methods like NMR and simulations, enabling cost-efficient, rapid, and accurate resolution of RNA conformational ensembles and also capturing rare excited and transient states which are critical for function yet elude experimental and computational detection due to their low populations or short lifetimes. Future applications of DynaRNA will range from contributing to current molecular dynamics simulation, RNA enhance sampling, interpreting RNA experiments, RNA-targeted drug binding site identification, RNA-protein binding mechanism. These capabilities make DynaRNA a powerful tool for paving the way for future advancements in RNA-targeted drug discovery and RNA therapy development such as mRNA vaccine design. Last but not least, DynaRNA breaks through the static structure prediction paradigm exemplified by AlphaFold3, pioneering the generative modeling of RNA dynamic ensembles—a framework that inherently aligns with RNA’s flexible nature, where biological functions emerge from continuous conformational transitions. By bridging RNA structure and dynamics, DynaRNA offers a scalable foundation for decoding the full complexity of RNA’s dynamic universe.

## Materials and Methods

### Dataset

DynaRNA was trained on high-quality RNA PDB crystal structures. We extracted 14,632 experimentally determined 3D RNA structures from RNAsolo database^41^. Training dataset was curated by removing entries that included non-RNA elements (e.g., DNAs and proteins), non-standard RNA elements (modified bases) or incomplete nucleotides. Structures containing 5 to 200 nucleotides were retained, producing 6,820 curated structures as the final training dataset. For test dataset, MD trajectories of five tetranucleotides were derived from previous REST2 simulations^10–11^, which were extensively sampled starting from both the experimental A-form conformations and the intercalated conformations.

### Formulation of DynaRNA

DynaRNA takes a single RNA structure as input, and does not rely on sequence features like MSA. Each nucleotide is coarse-grained into one particle located at C4’ atom, providing a minimal yet informative encoding of the RNA backbone geometry. The resulting representations are subsequently integrated into downstream modeling pipelines to facilitate structural learning. Denoising Diffusion Probabilistic Models (DDPM)^43^ is utilized in RNA conformation generation, which can be partitioned into a forward noising process and a symmetric backward denoising process. Both process are defined on discrete time space. The forward process gradually perturbs the original data with Gaussian noise, which is define by the following Itô stochastic differential equation (SDE):

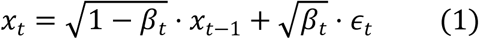

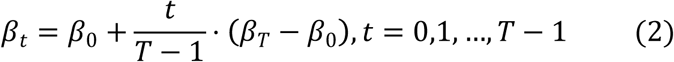

where *x_t_* represents the noised data at the *t-th* step, *β_t_* represents the noise level at step *t* which is defined by equation (2), *β*_0_ is set as 0.0001, *β_T_* is set as 0.02, *ɛ_t_∼* 𝒩(0, *I*) is Gaussian noise.

To reverse the noising process and recover original structures, we train an Equivariant Graph Neural Network (EGNN)^44^ to predict the noise *ɛ_θ_* (*x_t_*, *t*) given data *x_t_* at the time step *t*. The EGNN architecture is specifically designed to respect geometric symmetries such as translation and rotation equivariance, making it highly suitable for molecular or geometric data. Each layer of EGNN incorporates both node and edge updates to capture intricate geometric relationships among nucleotides. The node features consist of the 3D positions of C4’ atoms along with time-step embeddings, while edge features encode both the molecular connectivity and spatial distances. We used a hidden dimension of 128 across all layers, with LayerNorm and SiLU activation functions to enhance training stability and non-linearity. Temporal information is encoded using sinusoidal embeddings, following the standard approach in diffusion models. To reduce overfitting, dropout with a rate of 0.1 is applied after each EGNN layer. The final output of the network predicts the noise vector added at each time step, conditioned on both geometry and graph topology. During inference, the reverse (denoising) process is approximated by integrating the following formulation:

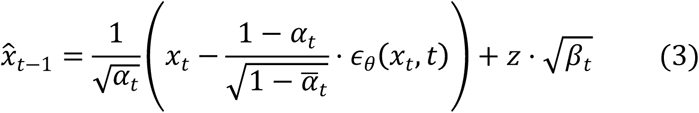

where *x̂_t_*_−1_ represents predicted data at the previous timestep *t* − 1, α*_t_* = 1 −*β_t_* represents the signal retention rate, 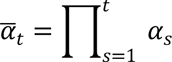 represents the cumulative signalretention, *ɛ_θ_* (*x_t_*, *t*) represents the noise predicted by EGNN, *z* ∼ 𝒩(0, *I*) represents the fresh Gaussian noise used during sampling.

In DDPM framework pursues a distinct training objective compared to other neural networks. Instead of directly fitting RNA coordinates, the network is designed to estimate noise in the perturbed data. We utilized L2 loss on noise defined as follows:

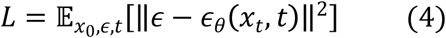

where *ɛ*represents the real noise, *ɛ_θ_* (*x_t_*, *t*) represents the noise predicted *ɛ_θ_* (*x_t_*, *t*) given data *x_t_* at the time step *t*. 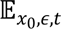 is used to calculate the mean square error (MSE) between them.

DynaRNA was implemented utilizing PyTorch and PyTorch-Lightning. All training processes were conducted on one NVIDIA 4090D GPU, taking approximately 14 days. The Model parameters were optimized with the Adam optimizer^45^, using a learning rate of 0.0001. To prevent gradient explosion and maintain numerical stability, gradient clipping with a maximum norm of 1.0 is applied. A weight decay of 1e-4 serves as a regularization mechanism to mitigate overfitting. Training procedure loss was shown in Figure S7. In our implementation of DynaRNA, we employed partial noising and denoising instead of conventional fully noising process. This approach aimed to balance efficient sampling and the preservation of essential initial structural information of RNA conformation generation. We implemented a partial noising process where the forward diffusion process as well as backward denoising process spans 800 steps instead of the full 1024 steps in inference.

### Analysis tools

Pymol was used for visualization and alignment. DSSR software was used for RNA structure analysis. Intercalation conformation was defined with nucleotide j is positioned between nucleotides i and i + 1 and forms stacking interactions with them, where j < i or j > i + 1. DBSCAN algorithm was used for clustering based on the RMSD of all heavy atoms where epsilon was set as 1.2 and min-points number was set as 10. GS and ES2 conformations were obtained from the PDB(8U3M). Initial structures of de novo folding tetraloop were constructed with the NAB module producing fully extended single-stranded conformations devoid of base pairing. Experimental structures from the PDB (2KOC and 8CLR) served as reference models for tetraloops.

## Data Availability

The Supplementary Information is still in progress, the dataset and code of DynaRNA will be available at GitHub.

## Acknowledgements

This work was supported by Shanghai Municipal Science and Technology Major Project, partially by SJTU Kunpeng & Ascend Center of Excellence, the Center for HPC at Shanghai Jiao Tong University, and the National Key Research and Development Program of China (2023YFF1205102 and 2020YFA0907700), the Fundamental Research Funds for the Central Universities (YG2023LC03), and the National Natural Science Foundation of China (32171242).

## Notes

### Competing Interest Statement

The authors have declared no competing interest.

## References

(1) Esteller, M. Non-coding RNAs in human disease. Nature Reviews Genetics 2011, 12 (12), 861–874.

(2) Lee, Y.-T.; Degenhardt, M. F. S.; Skeparnias, I.; Degenhardt, H. F.; Bhandari, Y. R.; Yu, P.; Stagno, J. R.; Fan, L.; Zhang, J.; Wang, Y.-X. The conformational space of RNase P RNA in solution. Nature 2025, 637 (8048), 1244–1251.

(3) Childs-Disney, J. L.; Yang, X.; Gibaut, Q. M. R.; Tong, Y.; Batey, R. T.; Disney, M. D. Targeting RNA structures with small molecules. Nature Reviews Drug Discovery 2022, 21 (10), 736–762.

(4) Bonilla, S. L.; Jang, K. Challenges, advances, and opportunities in RNA structural biology by Cryo-EM. Current Opinion in Structural Biology 2024, 88, 102894.

(5) Šponer, J.; Bussi, G.; Krepl, M.; Banáš, P.; Bottaro, S.; Cunha, R. A.; Gil-Ley, A.; Pinamonti, G.; Poblete, S.; Jurečka, P.;, et al. RNA Structural Dynamics As Captured by Molecular Simulations: A Comprehensive Overview. Chem Rev 2018, 118 (8), 4177–4338.

(6) Zhang, J.; Fei, Y.; Sun, L.; Zhang, Q. C. Advances and opportunities in RNA structure experimental determination and computational modeling. Nat Methods 2022, 19 (10), 1193–1207.

(7) Jones, D.; Allen, J. E.; Yang, Y.; Drew Bennett, W. F.; Gokhale, M.; Moshiri, N.; Rosing, T. S. Accelerators for Classical Molecular Dynamics Simulations of Biomolecules. Journal of Chemical Theory and Computation 2022, 18 (7), 4047–4069.

(8) Mlýnský, V.; Kührová, P.; Kühr, T.; Otyepka, M.; Bussi, G.; Banáš, P.; Šponer, J. Fine-Tuning of the AMBER RNA Force Field with a New Term Adjusting Interactions of Terminal Nucleotides. J Chem Theory Comput 2020, 16 (6), 3936–3946.

(9) Kührová, P.; Mlýnský, V.; Zgarbová, M.; Krepl, M.; Bussi, G.; Best, R. B.; Otyepka, M.; Šponer, J.; Banáš, P. Improving the Performance of the Amber RNA Force Field by Tuning the Hydrogen-Bonding Interactions. J Chem Theory Comput 2019, 15 (5), 3288–3305.

(10) Li, Z.; Song, G.; Zhu, J.; Mu, J.; Sun, Y.; Hong, X.; Choi, T.; Cui, X.; Chen, H. F. Excited-Ground-State Transition of the RNA Strand Slippage Mechanism Captured by the Base-Specific Force Field. J Chem Theory Comput 2024, 20 (14), 6082–6097.

(11) Li, Z.; Mu, J.; Chen, J.; Chen, H. F. Base-specific RNA force field improving the dynamics conformation of nucleotide. Int J Biol Macromol 2022, 222 (Pt A), 680–690.

(12) Subramaniam, S. Structural biology in the age of AI. Nature Methods 2024, 21 (1), 18–19.

(13) Jumper, J.; Evans, R.; Pritzel, A.; Green, T.; Figurnov, M.; Ronneberger, O.; Tunyasuvunakool, K.; Bates, R.; Žídek, A.; Potapenko, A.;, et al. Highly accurate protein structure prediction with AlphaFold. Nature 2021, 596 (7873), 583–589.

(14) Abramson, J.; Adler, J.; Dunger, J.; Evans, R.; Green, T.; Pritzel, A.; Ronneberger, O.; Willmore, L.; Ballard, A. J.; Bambrick, J.;, et al. Accurate structure prediction of biomolecular interactions with AlphaFold 3. Nature 2024, 630 (8016), 493–500.

(15) Ding, J.; Lee, Y.-T.; Bhandari, Y.; Schwieters, C. D.; Fan, L.; Yu, P.; Tarosov, S. G.; Stagno, J. R.; Ma, B.; Nussinov, R.;, et al. Visualizing RNA conformational and architectural heterogeneity in solution. Nature Communications 2023, 14 (1), 714.

(16) Zhu, J.; Li, Z.; Zhang, B.; Zheng, Z.; Zhong, B.; Bai, J.; Wang, T.; Wei, T.; Yang, J.; Chen, H.-F. Precise Generation of Conformational Ensembles for Intrinsically Disordered Proteins Using Fine-tuned Diffusion Models. bioRxiv 2024, 2024.2005.2005.592611.

(17) Condon, D. E.; Kennedy, S. D.; Mort, B. C.; Kierzek, R.; Yildirim, I.; Turner, D. H. Stacking in RNA: NMR of Four Tetramers Benchmark Molecular Dynamics. Journal of Chemical Theory and Computation 2015, 11 (6), 2729–2742.

(18) Nozinovic, S.; Fürtig, B.; Jonker, H. R.; Richter, C.; Schwalbe, H. High-resolution NMR structure of an RNA model system: the 14-mer cUUCGg tetraloop hairpin RNA. Nucleic Acids Res 2010, 38 (2), 683–694.

(19) Oxenfarth, A.; Kümmerer, F.; Bottaro, S.; Schnieders, R.; Pinter, G.; Jonker, H. R. A.; Fürtig, B.; Richter, C.; Blackledge, M.; Lindorff-Larsen, K.;, et al. Integrated NMR/Molecular Dynamics Determination of the Ensemble Conformation of a Thermodynamically Stable CUUG RNA Tetraloop. J Am Chem Soc 2023, 145 (30), 16557–16572.

(20) Roy, R.; Geng, A.; Shi, H.; Merriman, D. K.; Dethoff, E. A.; Salmon, L.; Al-Hashimi, H. M. Kinetic Resolution of the Atomic 3D Structures Formed by Ground and Excited Conformational States in an RNA Dynamic Ensemble. J Am Chem Soc 2023, 145 (42), 22964–22978.

(21) Geng, A.; Ganser, L.; Roy, R.; Shi, H.; Pratihar, S.; Case, D. A.; Al-Hashimi, H. M. An RNA excited conformational state at atomic resolution. Nat Commun 2023, 14 (1), 8432.

(22) Trippe, B. L.; Yim, J.; Tischer, D.; Baker, D.; Broderick, T.; Barzilay, R.; Jaakkola, T. Diffusion probabilistic modeling of protein backbones in 3d for the motif-scaffolding problem. *arXiv preprint arXiv:2206.04119* 2022.

(23) Lu, J.; Zhong, B.; Zhang, Z.; Tang, J. Str2str: A score-based framework for zero-shot protein conformation sampling. *arXiv preprint arXiv:2306.03117 2023.*

(24) Batzner, S.; Musaelian, A.; Sun, L.; Geiger, M.; Mailoa, J. P.; Kornbluth, M.; Molinari, N.; Smidt, T. E.; Kozinsky, B. E(3)-equivariant graph neural networks for data-efficient and accurate interatomic potentials. Nature Communications 2022, 13 (1), 2453.

(25) Soleymani, F.; Paquet, E.; Viktor, H. L.; Michalowski, W. Structure-based protein and small molecule generation using EGNN and diffusion models: A comprehensive review. Comput Struct Biotechnol J 2024, 23, 2779–2797.

(26) Burley, S. K.; Bhatt, R.; Bhikadiya, C.; Bi, C.; Biester, A.; Biswas, P.; Bittrich, S.; Blaumann, S.; Brown, R.; Chao, H.;, et al. Updated resources for exploring experimentally-determined PDB structures and Computed Structure Models at the RCSB Protein Data Bank. Nucleic Acids Res 2025, 53 (D1), D564–d574.

(27) Zgarbová, M.; Otyepka, M.; Sponer, J.; Mládek, A.; Banáš, P.; Cheatham, T. E., 3rd; Jurečka, P. Refinement of the Cornell et al. Nucleic Acids Force Field Based on Reference Quantum Chemical Calculations of Glycosidic Torsion Profiles. J Chem Theory Comput 2011, 7 (9), 2886–2902.

(28) Bannwarth, S.; Gatignol, A. HIV-1 TAR RNA: the target of molecular interactions between the virus and its host. Curr HIV Res 2005, 3 (1), 61–71.

(29) Bou-Nader, C.; Link, K. A.; Suddala, K. C.; Knutson, J. R.; Zhang, J. Structures of complete HIV-1 TAR RNA portray a dynamic platform poised for protein binding and structural remodeling. Nature Communications 2025, 16 (1), 2252.

(30) Chu, C. C.; Plangger, R.; Kreutz, C.; Al-Hashimi, H. M. Dynamic ensemble of HIV-1 RRE stem IIB reveals non-native conformations that disrupt the Rev-binding site. Nucleic Acids Res 2019, 47 (13), 7105–7117.

(31) Xue, Y.; Kellogg, D.; Kimsey, I. J.; Sathyamoorthy, B.; Stein, Z. W.; McBrairty, M.; Al-Hashimi, H. M. Characterizing RNA Excited States Using NMR Relaxation Dispersion. Methods Enzymol 2015, 558, 39–73.

(32) Han, G.; Xue, Y. Rational design of hairpin RNA excited states reveals multi-step transitions. Nature Communications 2022, 13 (1), 1523.

(33) Klosterman, P. S.; Hendrix, D. K.; Tamura, M.; Holbrook, S. R.; Brenner, S. E. Three-dimensional motifs from the SCOR, structural classification of RNA database: extruded strands, base triples, tetraloops and U-turns. Nucleic Acids Res 2004, 32 (8), 2342–2352.

(34) Thapar, R.; Denmon, A. P.; Nikonowicz, E. P. Recognition modes of RNA tetraloops and tetraloop-like motifs by RNA-binding proteins. Wiley Interdiscip Rev RNA 2014, 5 (1), 49–67.

(35) Tinoco, I.; Bustamante, C. How RNA folds. Journal of Molecular Biology 1999, 293 (2), 271–281.

(36) Chen, A. A.; García, A. E. High-resolution reversible folding of hyperstable RNA tetraloops using molecular dynamics simulations. Proceedings of the National Academy of Sciences 2013, 110 (42), 16820–16825.

(37) Kührová, P.; Best, R. B.; Bottaro, S.; Bussi, G.; Šponer, J.; Otyepka, M.; Banáš, P. Computer Folding of RNA Tetraloops: Identification of Key Force Field Deficiencies. Journal of Chemical Theory and Computation 2016, 12 (9), 4534–4548.

(38) Mustoe, A. M.; Brooks, C. L.; Al-Hashimi, H. M. Hierarchy of RNA functional dynamics. Annu Rev Biochem 2014, 83, 441–466.

(39) Ganser, L. R.; Kelly, M. L.; Herschlag, D.; Al-Hashimi, H. M. The roles of structural dynamics in the cellular functions of RNAs. Nat Rev Mol Cell Biol 2019, 20 (8), 474–489.

(40) Duzdevich, D.; Redding, S.; Greene, E. C. DNA Dynamics and Single-Molecule Biology. Chemical Reviews 2014, 114 (6), 3072–3086.

(41) Adamczyk, B.; Antczak, M.; Szachniuk, M. RNAsolo: a repository of cleaned PDB-derived RNA 3D structures. Bioinformatics 2022, 38 (14), 3668–3670.

(42) Song, Y.; Sohl-Dickstein, J.; Kingma, D. P.; Kumar, A.; Ermon, S.; Poole, B. Score-based generative modeling through stochastic differential equations. *arXiv preprint arXiv:2011.13456 2020.*

(43) Ho, J.; Jain, A.; Abbeel, P. Denoising diffusion probabilistic models. Advances in neural information processing systems 2020, 33, 6840–6851.

(44) Satorras, V. G.; Hoogeboom, E.; Welling, M. E (n) equivariant graph neural networks. In International conference on machine learning, 2021; PMLR: pp 9323–9332.

(45) Kingma, D. P.; Ba, J. Adam: A method for stochastic optimization. *arXiv preprint arXiv:1412*.6980 2014.

