## Supplemental Information for "DynaRNA: Dynamic RNA Conformation Ensemble Generation with Diffusion Model"

*Corresponding Author

Hai-Feng Chen (Full Professor)

Notes

The authors declare that there is no conflict of interest.

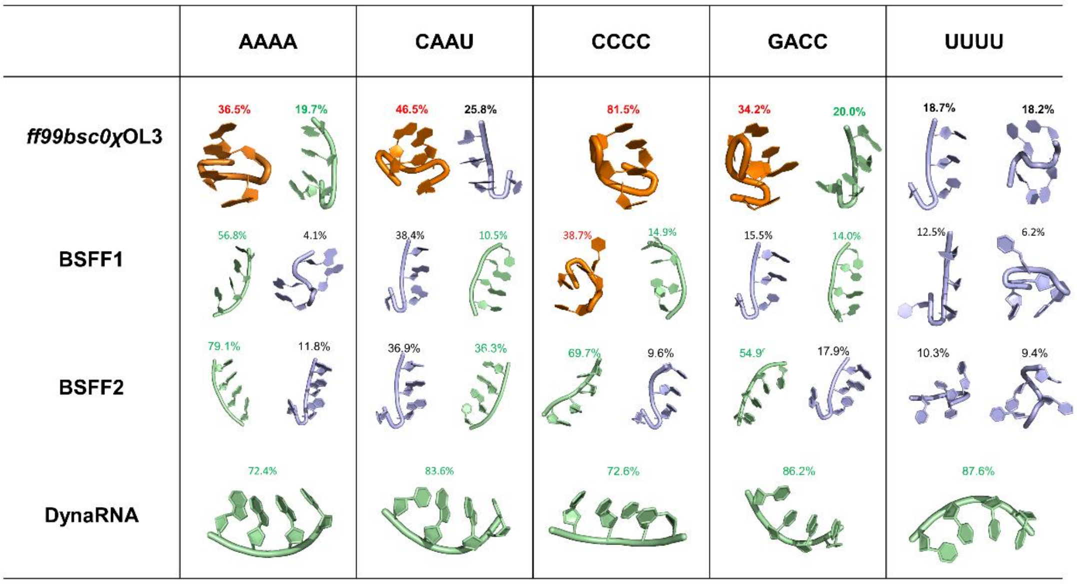


**Fig S1.** Clustering results of tetranucleotides conformation ensembles generated with molecular dynamics simulations with OL3, BSFF1, BSFF2, and DynaRNA initialized from A-form structures. The orange color represents the intercalated conformation, the green color represents the A-form conformation, the blue color represents the other conformations.

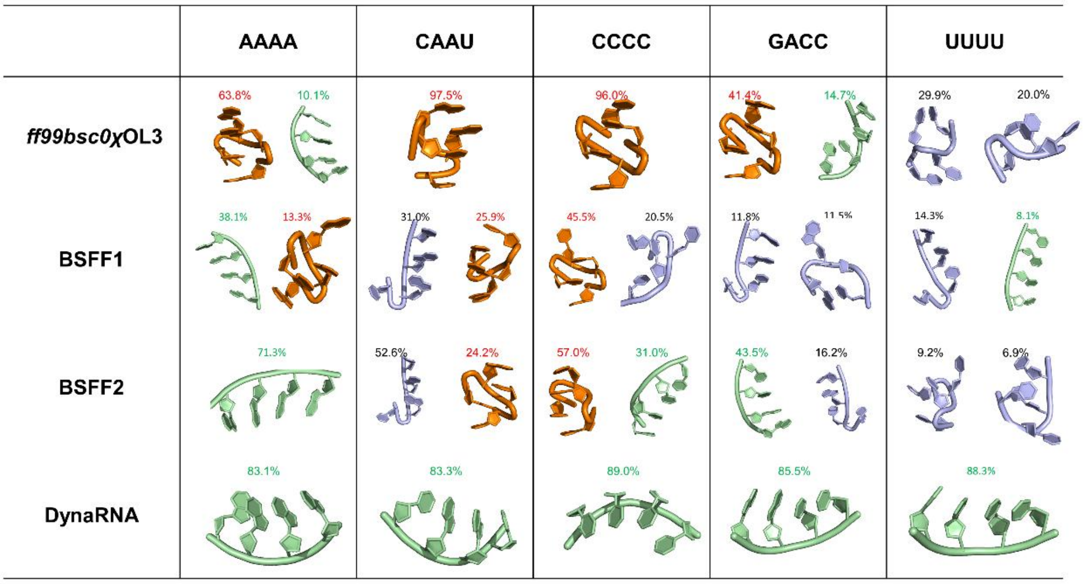


**Fig S2.** Clustering results of tetranucleotides conformation ensembles generated with molecular dynamics simulations with OL3, BSFF1, BSFF2, and DynaRNA initialized from A-form structures. The orange color represents the intercalated conformation, the green color represents the A-form conformation, the blue color represents the other conformations.

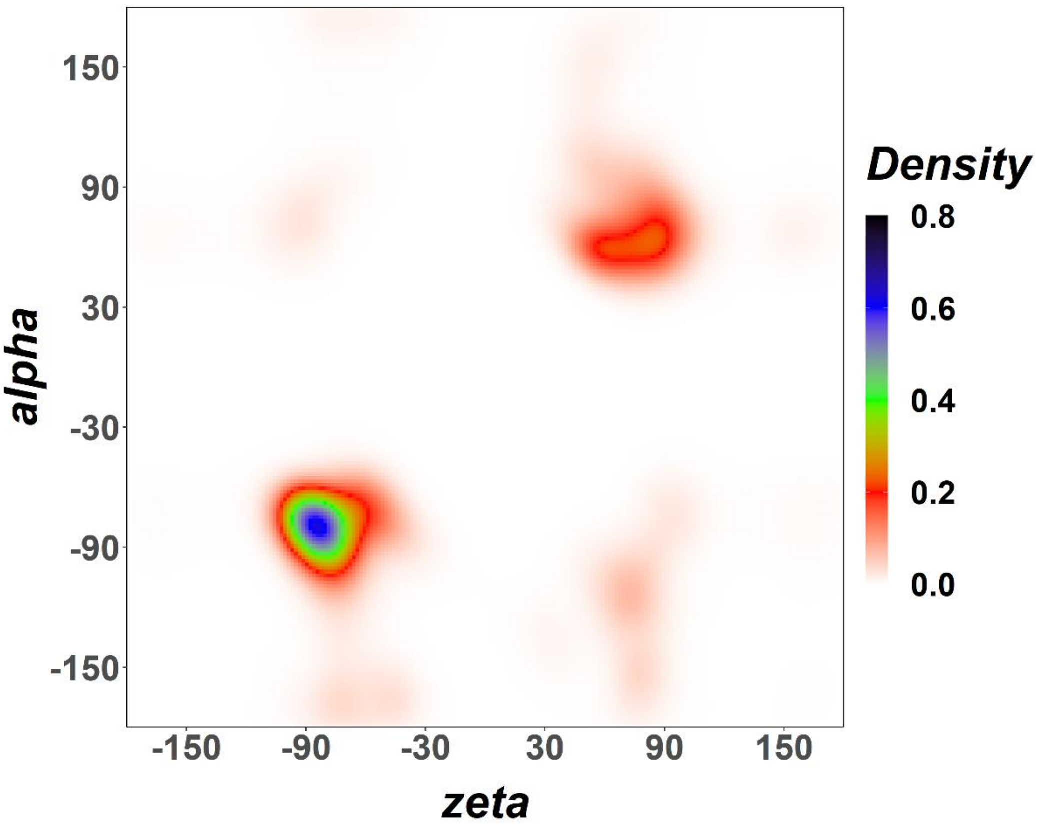


**Figure S3.** Dihedral statistical distribution of conformation ensemble generated with molecular dynamics simulations with BSFF1 initialized from A-form structures.

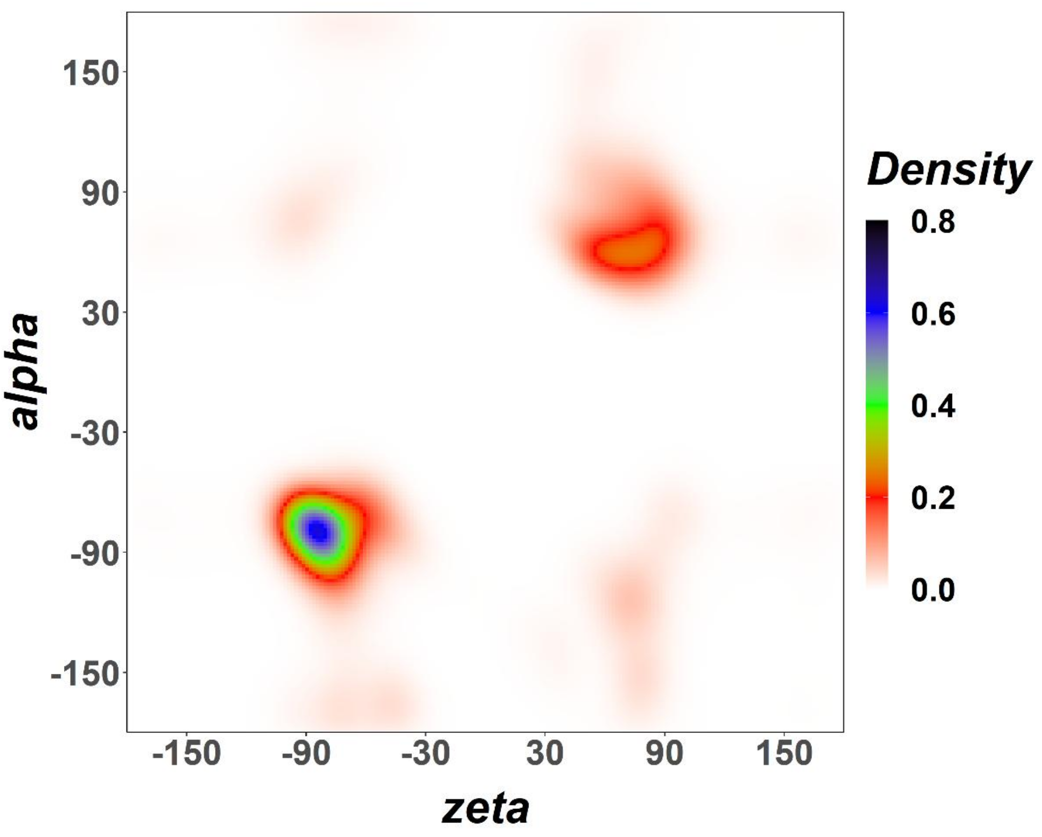


**Figure S4.** Dihedral statistical distribution of conformation ensemble generated with molecular dynamics simulations with BSFF1 initialized from intercalated structures.

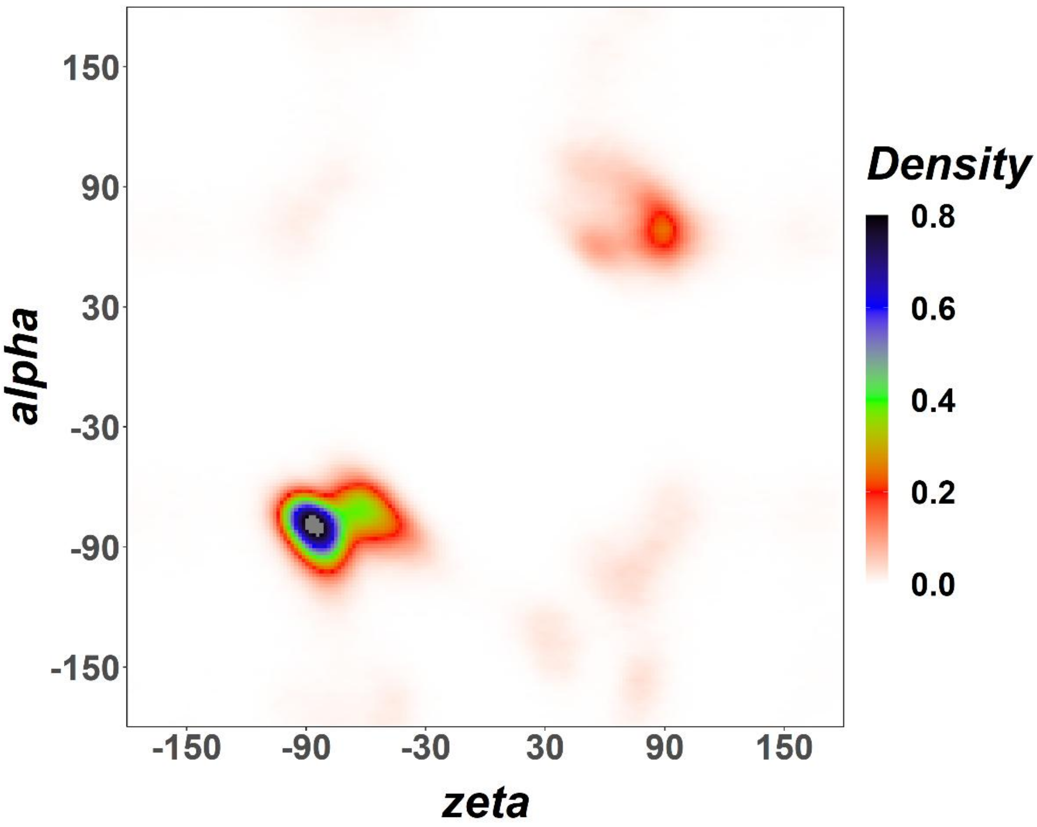


**Figure S5.** Dihedral statistical distribution of conformation ensemble generated with molecular dynamics simulations with BSFF2 initialized from A-form structures.

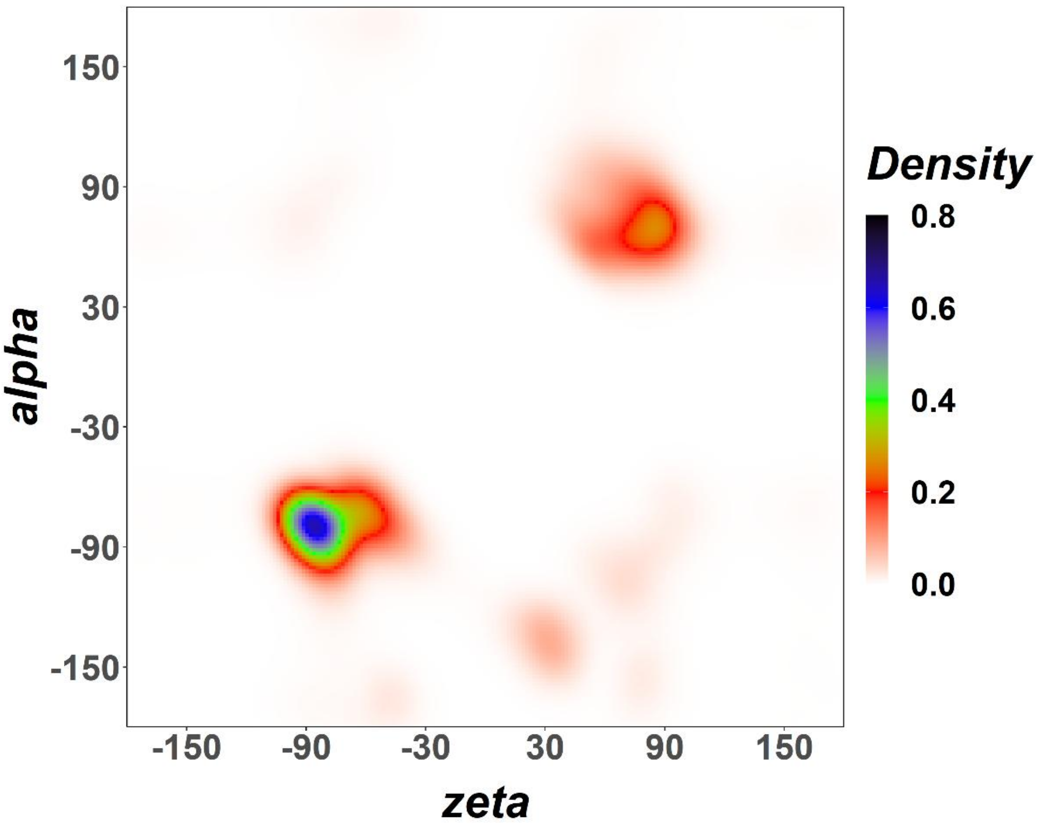


**Figure S6.** Dihedral statistical distribution of conformation ensemble generated with molecular dynamics simulations with BSFF2 initialized from intercalated structures.

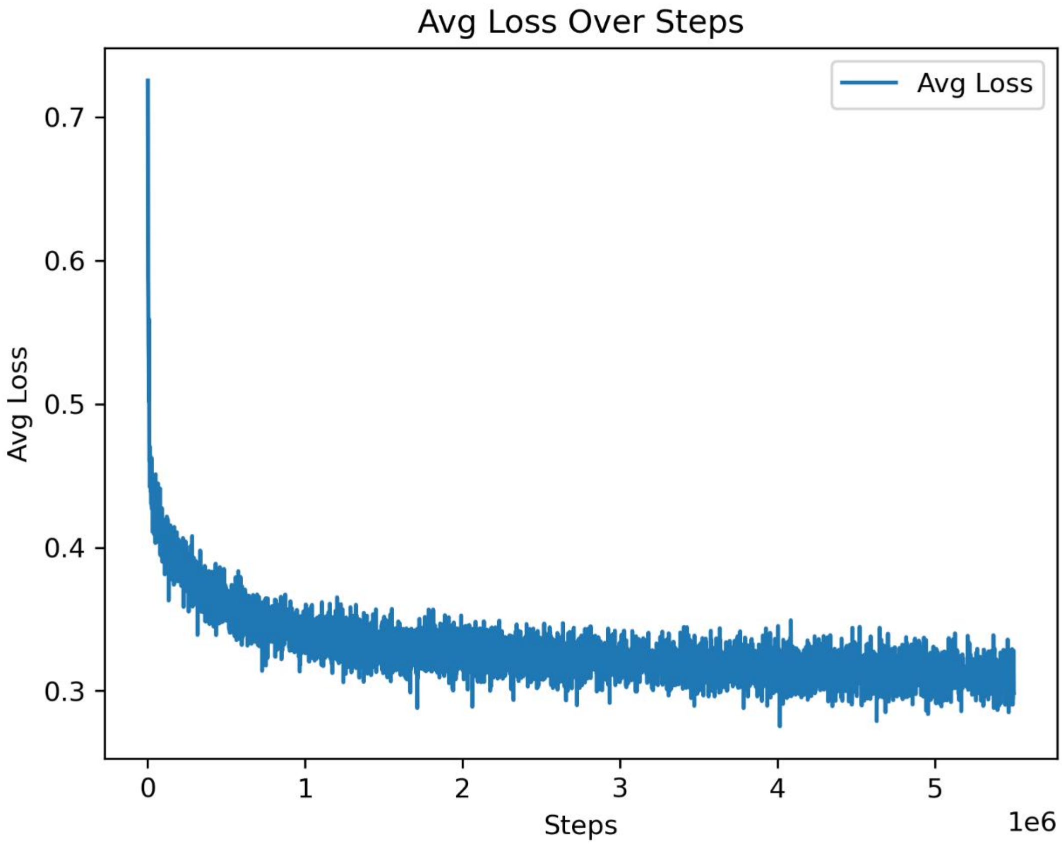


Fig S7. Loss curves of model over training steps.
